# A simple, inexpensive battery-powered homeothermic warming pad for mice and rats

**DOI:** 10.1101/2025.04.28.651075

**Authors:** Isabella R. Fleites, Kevin Morales, Stephen D. Roper

## Abstract

**Background:** Anesthesia decreases core body temperature, and this seriously compromises the physiological status of an experimental animal. Hypothermia alters many aspects of neural function. When recording nervous system activity in anesthetized animals, their core temperature must be stabilized.

**New method:** This report describes an inexpensive, battery-powered, temperature-controlled warming pad for mice and rats and documents its validity and utility. The device is portable, making it convenient for researchers who conduct procedures such as surgical preparations in one location and transport the anesthetized animal to another location for experimental recordings.

**Results:** The device keeps anesthetized mice normothermic ±0.7° for over 6 hours without supplemental warmth (e.g., heat lamp), despite >15° differential between ambient room temperature and core body temperature. We demonstrate how the warming pad can be used for *in vivo* imaging of neuronal activity for a prolonged period in mice.

**Comparison with existing methods:** Commercial heating pads for small animals are expensive, somewhat bulky, and require power cords and a 120/240V source. Transporting an anesthetized animal from one location (e.g. surgical suite) to another (e.g., imaging rig) involves moving power cords. Moreover, commercial devices are not always compatible with custom stereotaxic frames, microscope stages, or holding boxes. The device described here is small, inexpensive, battery-powered, and readily adaptable to experimental set ups.

**Conclusion:** The homeothermic heating pad provides a simple method for maintaining the core temperature of anesthetized small animals. It can be constructed in under 30 minutes, the components are readily available, and the cost is less than $100. It is exceptionally useful for experiments on mice or rats.

**Highlights:**

- Temperature-controlled warming pad for small rodents
- Low cost, well under $100
- Components available from online suppliers
- Can be assembled in less than 30 minutes
- Does not require specialty tools
- The device is stable and robust

## Introduction

Present day methods to investigate neural activity *in vivo* in brains of mice and rats utilize sophisticated functional imaging techniques and/or multielectrode arrays [1-4]. In many such studies, the experimental animal is deeply sedated or in a surgical plane of anesthesia. Yet, it is well recognized that anesthesia incapacitates thermoregulation in animals; the core temperature of an anesthetized animal rapidly drops. For example, in rats and mice intraperitoneal injection of a commonly-used anesthetic, ketamine, quickly leads to hypothermia [5-7]. Indeed, hypothermia complicates general anesthesia in humans and rodents alike [8-11]. Consequently, a device to maintain an animal’s core body temperature is required for a wide range of experimentation on anesthetized animals [e.g. [12-18].

Hypothermia is a particular problem in neuroscience. Low temperature affects brain and neuronal function in a number of ways. Mild hypothermia during anesthesia strengthens the blood brain barrier against experimental insults such as perfusion with hyperosmolar solutions [19]; more severe hypothermia disrupts the blood brain barrier [20]. Hypothermia reduces nerve conduction velocity in the central [21] and peripheral [22] nerves and alters many aspects synaptic transmission [23, 24](*cf*. in cold-blooded animals [25]). Clearly, maintaining the core body temperature of an experimental animal is critical when studying neuronal activity, emphasizing the importance of some form of warming device, especially for anesthetized animals.

There are several commercially available warming pads for small animals, many of which include controllers with temperature probe feedback to maintain the animal at a constant core temperature [26-32]. These commercial devices are dependable and stable, yet often rather bulky and somewhat costly. Other, more specialized homeothermic heating units for rodents have been described but have limited applicability for general use [33-37]. Moreover, commonly available homeothermic heating devices have power supplies that must be plugged into a 120 or 240 V outlet. These factors can be limiting. The requirement for an 120/240 V power can be especially vexing if an animal is surgically prepared for experimentation at a different location from where the actual recording takes place. Transporting the preparation, even a short distance, requires moving a controller/power supply and power cord. Moreover, power supplies/controllers for warming pads can introduce noise in electrophysiological recordings.

In our studies using *in vivo* functional imaging of sensory neuron activity in mice, we have overcome these aggravations by designing an extremely inexpensive, easy-to-construct, and supremely portable homeothermic warming pad. This warming pad is powered by a small lithium polymer (LiPo) battery such as those used to power drones. This device requires minimal skills to construct. All parts are off-the-shelf items, readily available by online order. The only tools needed are a small screwdriver, a soldering iron, and a wire-cutter/stripper. The total cost is well below $100, and the entire construction takes less than 30 minutes. The small size and flexible arrangement of its components makes the homeothermic warming pad readily adaptable to a custom stereotaxic frame, recording platforms, or observation cages, or as in our case, the stage of an upright confocal microscope.

Although we use this homeothermic warming pad for *in vivo* calcium imaging of neuronal activity in surgically-exposed cranial sensory ganglia [38, 39], any studies using sedated or anesthetized small rodents would benefit by this device. In our experience, when powered by fully charged LiPo batteries, the heating pad maintains the core body temperature of an anesthetized mouse stable more than 6 hours.

## 2. Materials and methods

### 2.1 Animals

We have used the battery-powered homeothermic heating pad on 34 mice over a span of 8 months. All experimental procedures were conducted according to the National Institutes of Health Guidelines for the Care and Use of Laboratory Animals and authorized by the University of Miami Institutional Animal Care and Use Committee. The animals are transgenic mice that express GCaMP6s in sensory ganglion neurons. [These mice were generated by crossing floxed GCaMP6s mice (stock no. 024106, Jackson Laboratory, Bar Harbor, ME, USA) with Pirt-Cre mice (obtained from X. Dong, Johns Hopkins)]. However, the specific strain of mice is not critical for the development, construction, or use of the battery-powered homeothermic heating pad described below. Details of the transgenic strain are included here only to support the imaging data we present illustrating how we use the heating pad. The device can be used for any small rodent.

### 2.2 *In vivo* confocal calcium imaging to illustrate one use of the warming pad

We tested and validated the homeothermic heating pad in rather stringent conditions. The device has been used for over ½ year for *in vivo* confocal calcium imaging of sensory ganglion neurons in anesthetized mice. These experiments are carried out in an air-conditioned room having an ambient temperature of 20° to 21°. Mice were deeply anesthetized with a mixture of Ketamine (80-100 mg/kg) and Xylazine (10 mg/kg), the brain exposed, and cortex overlying the trigeminal and geniculate ganglion aspirated to unilaterally expose the cranial ganglia for functional imaging. Details of these procedures can be found in references [38, 39]. The core temperature of the animals as measured by the device’s rectal temperature probe (bead thermistor) remained constant and the animal kept in stable physiological condition for over 6 hours using the homeothermic heating pad and no other auxiliary heating (*e*.*g*., infrared lamp, blanket).

### 2.3 Warming pad components

The warming pad consists of a small silicone rubber heating pad attached to a relay switch that is operated by a small commercially available temperature control circuit. A 7.6V LiPo battery powers the temperature control circuit as well as the silicone heating pad. A 10K bead thermistor with insulated leads serves as a rectal temperature probe and provides feedback for the temperature control circuit. The silicone pad and temperature control circuit draw less than 1 A during operation, well within the specifications for the silicone heating pad (15 W) and allowing all connections to be safely made with flexible 20 ga hookup wire.

The specific components, their sources, and their cost, are listed in Table 1.

**Table 1.** List of components to construct homeothermic warming pad

| component | num<br>ber | cost | source |
| --- | --- | --- | --- |
| 12V 15W 50 mm x 100 mm silicone heating pad | 1 | \$15.99<br>4/Pkg | Amazon<br>https://www.amazon.com/DERNORD-Silicone-Heating-Rubber-Flexible/dp/B0BDYBTNCD?ref_=ast_sto_dp&th=1 |
| Murata NTC BEAD Thermistor 10KOhm 3380K NXFT15XH103FA2B025 | 1 | \$ 0.60 | Digikey<br>https://www.digikey.com/en/products/detail/murata-electronics/NXFT15XH103FA2B025/2533819?s=N4lgTCBcDaIHIA0BiAVAjAVgQCTQBgGYkB BMAITzAxAF0BfIA |
| JST-XH 2.54mm 1S 2 Pin Balance Plug Lead Socket male connector | 1 | \$7.99<br>5/pkg | Amazon<br>https://www.amazon.com/dp/B07YKHV46N?psc=1&ref=ppx_yo2ov_dt_b_product_details |
| JST Plug Connector 2 Pin Male with 20 AWG leads | 1 | \$7.49<br>10/Pkg | Amazon<br>https://www.amazon.com/dp/B01M5AHF0Z?ref_=ppx_hzsearch_conn_dt_b_fed_asin_title_1 |
| 7.4V 1200mAh LiPo Battery | 1 | \$23.99<br>2/Pkg | Amazon<br>https://www.amazon.com/dp/B0B74LR9R7?ref_=ppx_hzsearch_conn_dt_b_fed_asin_title_1 |
| DROK Electronic Thermostat Controller | 1 | \$11.99 | Amazon<br>https://www.amazon.com/DROK-Electronic-Thermostat-Controller-Temperature/dp/B07GQPT9VG?ref_=ast_sto_dp&th=1 |

### 2.4 Warming pad construction

Lay out the items needed for construction (Fig. 1):

a. 12V 15W 50 mm x 100 mm silicone heating pad (50 mm x 100 mm is a convenient size for mice. Larger silicone rubber heating pads would be more appropriate for rats)
b. NTC Thermistor NXFT15XH103FA2B025
c. JST-XH 2.54mm 1S 2 Pin Balance Plug Lead Socket male connector
d. JST Plug Connector 2 Pin Male with 20 AWG leads
e. 7.4V 1200mAh LiPo Battery
f. temperature controller circuit
g. short length of 20 AWG hookup wire

**Figure 1.**
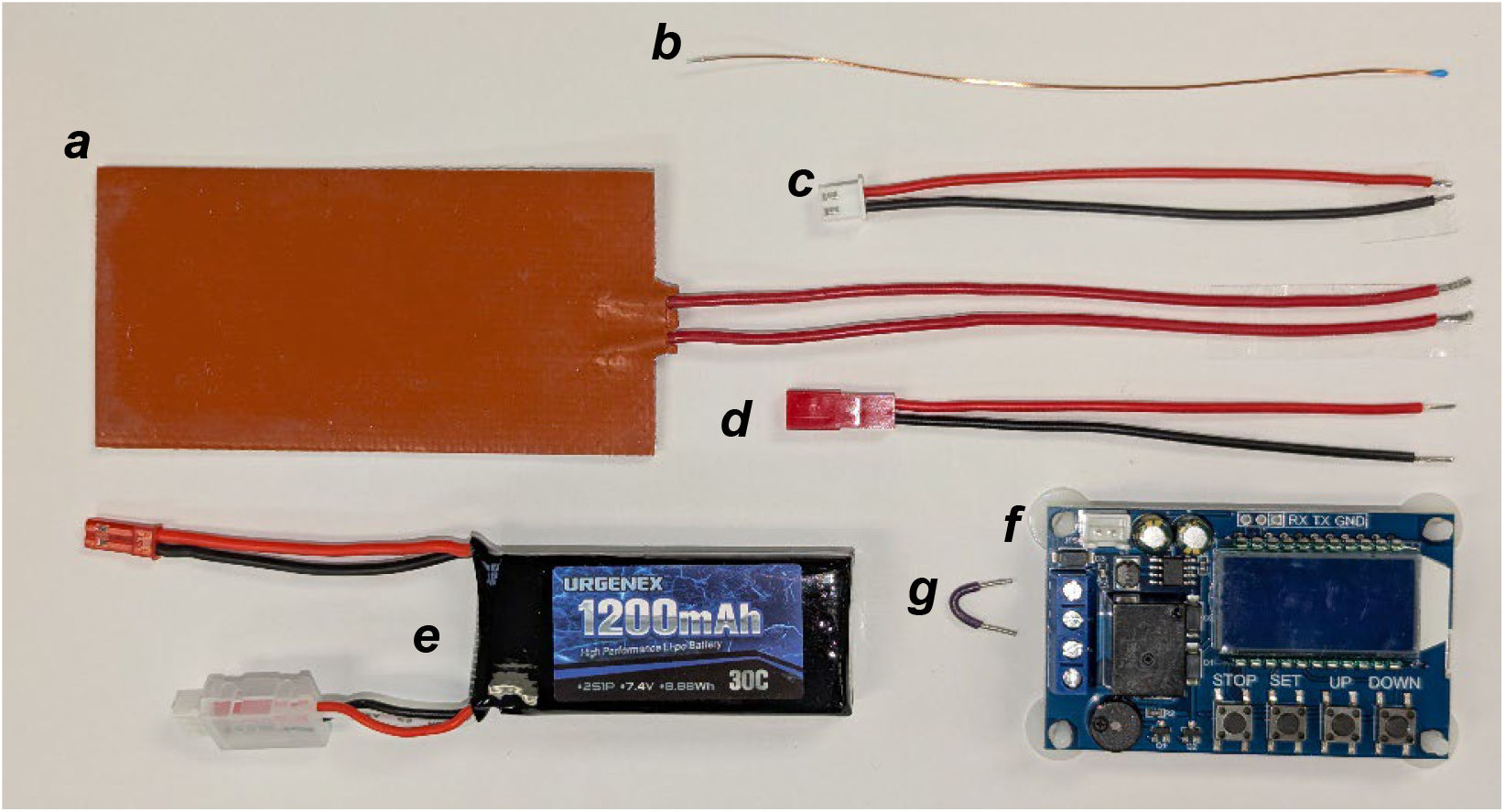
Layout of parts for homeothermic heating pad construction. ***a*** to **g**, see text.

Begin by inserting the two leads from the silicone heating pad (cf. Fig. 1*a*) into terminals #2 and #4 of the temperature control circuit (cf. Fig. 1*f*) and tighten the terminal screws as shown in Figure 2. Best practice is to tin the leads before attaching to control board terminal.

**Figure 2.**
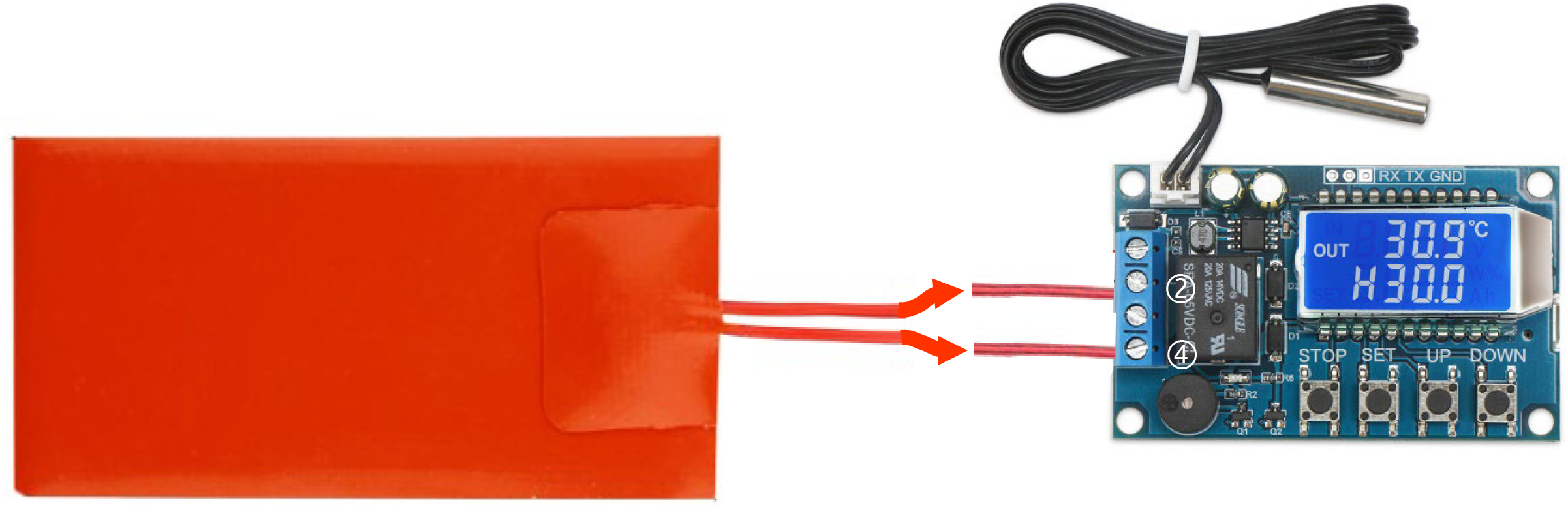
Attach silicone heating pad to temperature control circuit. Note, thermistor probe that comes with the temperature control circuit is shown here.

Next, attach the leads from the 2-pin male JST plug (cf. Fig. 1d) to the temperature control circuit as shown in Figure 3. Connect the red (positive) lead to terminal #1 and the black (negative) lead to terminal #2. Add a short 20 Ga hookup wire jumper between terminals #1 and #3 (Fig. 3). (Hint: you can salvage a short piece of hookup wire from the heating pad leads if you have cut those leads to a shorter length). As above, best practice is to tin all leads before clamping into control board terminal.

**Figure 3.**
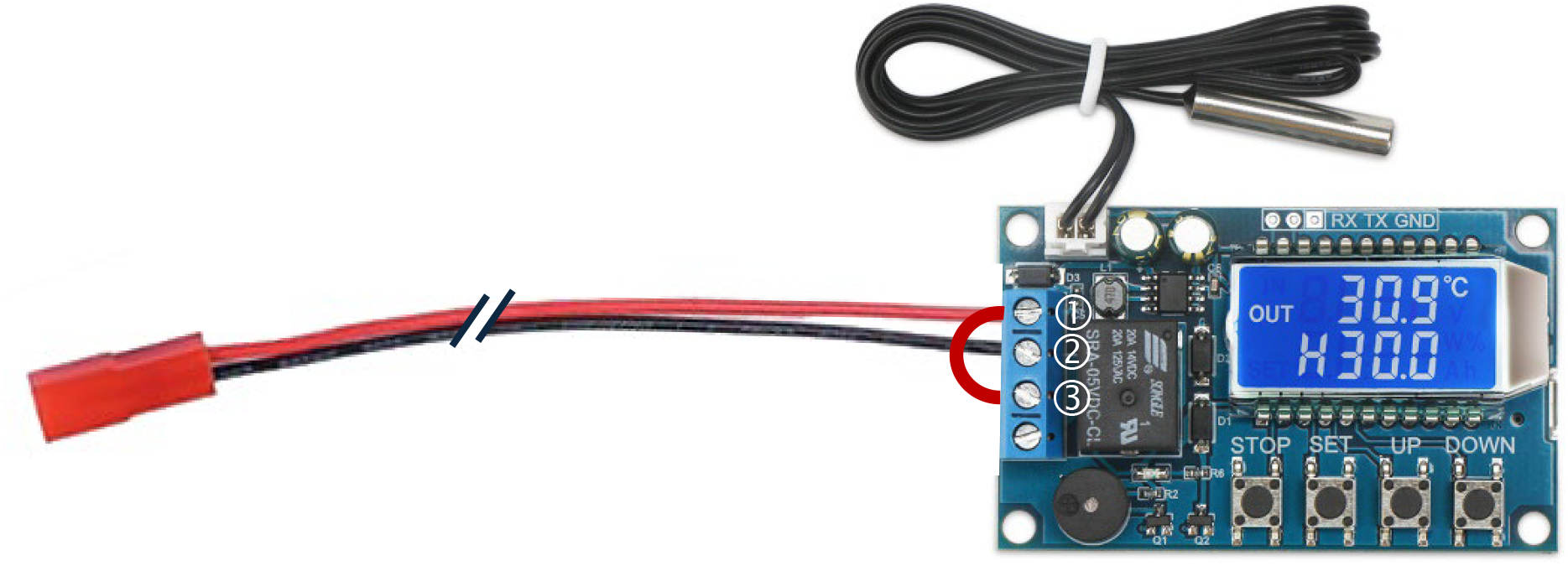
Attach leads from JST plug to terminals 1 and 2. Connect terminals 1 and 3 with a short hookup wire. Note, for clarity, silicone heating pad connections (Fig. 2) are not shown.

Finally, solder leads from JST-XH plug (cf. Fig. 1c) to the bead thermistor (cf. Fig. 1b). Remove (unplug) thermistor probe that is shipped with the temperature controller and replace with the bead thermistor probe as shown in Figure 4. The bead thermistor serves as a rectal thermal probe to monitor the core temperature of the mouse or rat.

**Figure 4.**
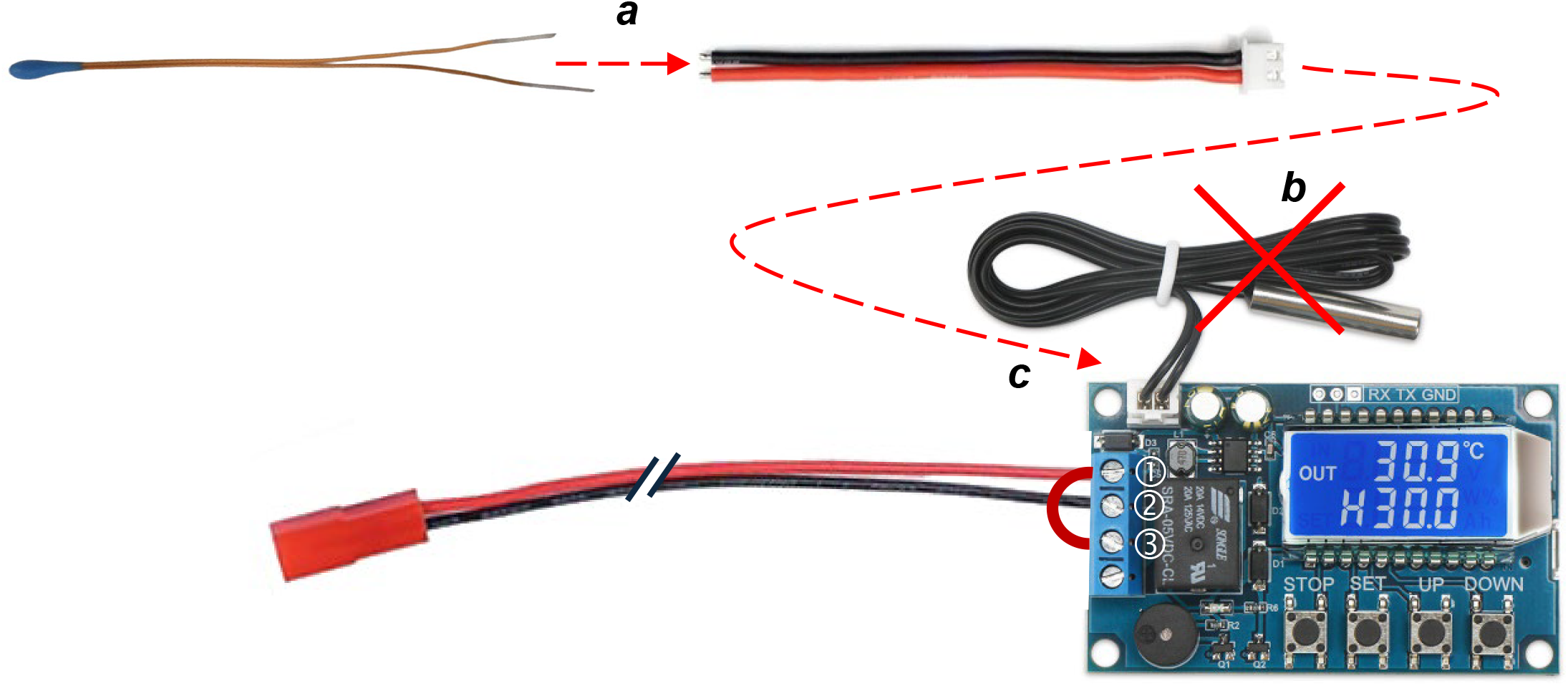
Prepare the rectal thermal probe. ***a***, Solder JST-HX plug to bead thermistor. ***b***, unplug the thermistor probe that comes with the temperature controller. ***c***, plug bead thermistor lead into controller. Note, for clarity, silicone heating pad connections (Fig. 2) are not shown.

This completes the construction (Fig. 5). The device is now ready to use and can be mounted on an appropriate platform for experimentation. It is convenient to mount the temperature controller circuit on tall nylon screws (e.g., 1” #6 screws) because the 7.4 V LiPo battery then fits neatly and conveniently underneath the circuit board. An example of the fully operational unit is shown in Figure 5.

**Figure 5.**
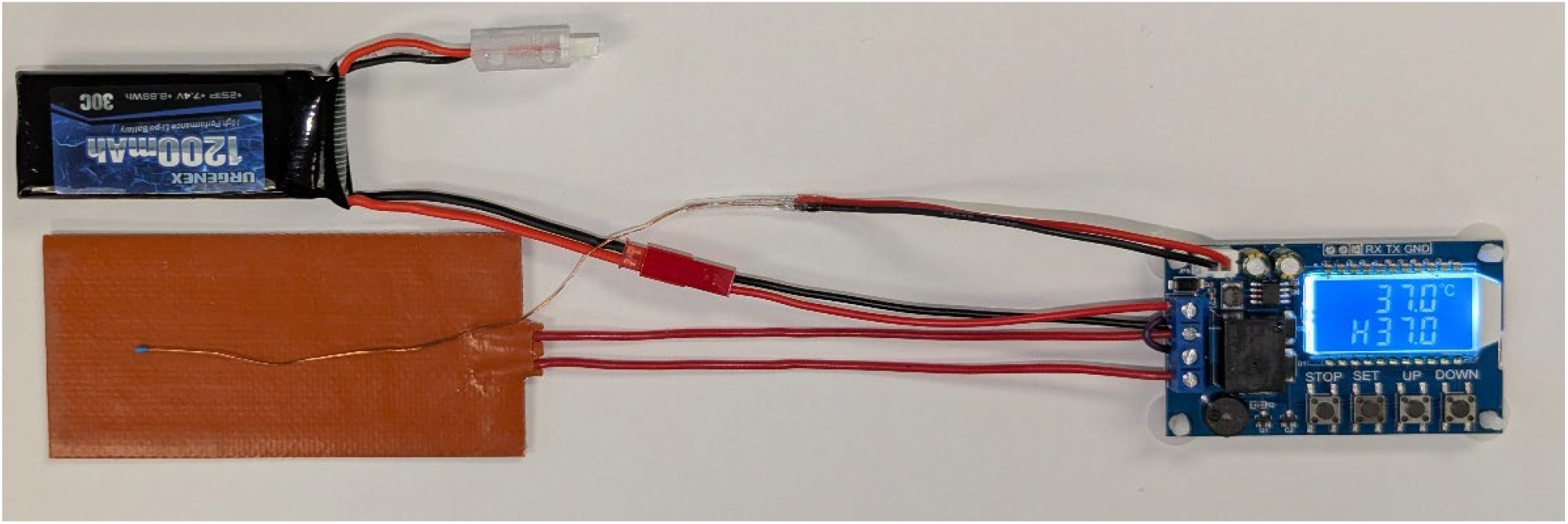
A fully completed homeothermic warming pad device, operating in a room with ambient temperature of 22°. The sensor probe is resting on the silicone rubber pad and shows the temperature at the surface of the warm pad (37°).

### 2.5. Operation instructions

To use the homeothermic heating pad, place a small amount of Vaseline on the bead thermometer and gently insert into rectum of anesthetized/sedated mouse or rat. Best practice is to tape the lead of the thermistor to the animal’s tail to prevent the probe from becoming dislodged. Place the animal prone onto the silicone heating pad. Power the device by plugging in the 7.4 V LiPo battery. Press the “Set” button on the thermostat controller. “H” (heating) will show next to the set temperature. Press “Set” once more to activate the temperature setting. Press “Up” or “Down” button to change the set temperature to your specifications. For example, use a set temperature of 37°. Hold the “Set” button for 3 seconds (or do not press any button for 6 sec) and device is now fully operational.

If the animal’s core temperature is lower than the set temperature (e.g., 37°), the relay powering the silicone heating pad will close. Current will be delivered to the silicone heating pad, the red light will come on, and the silicone pad will begin to warm up until the rectal probe exceeds the set temperature by a fraction of a degree (“hysteresis”; factory setting is 0.5°). When the rectal probe exceeds the set temperature plus hysteresis (in this example, 37.5°), the relay opens, the red light goes off, power is cut from the silicone heating pad, and the heating pad and the animal begin to cool (assuming an ambient room temperature < 37.5°). When the rectal probe drops to the set temperature minus hysteresis (e.g., 36.5°), the relay will close, and power will once again be delivered to the silicone heating pad to warm the animal. Hysteresis can be set by pressing the “Set” button three times and using “Up” or “Down” buttons. More detailed instructions for operating the temperature controller come with the device, including a serial interface control, should this be of interest to the users.

## 3. Results

In a critical analytical test of the device, we maintained a small (18.5 g) male mouse under deep anesthesia (as above) for 6 hours on the warming pad. We did not expose the brain but kept the anesthetized mouse in the open, without cover, in an air-conditioned room at an ambient temperature of 20.3°. Booster injections of ketamine were given as necessary to maintain a surgical plane of anesthesia. The mouse sustained a normal respiration rate for the entire 6 hours, varying from an estimated low of 110/min to a high of 215/min, depending on how long had elapsed from a ketamine booster injection (consistent with ketamine’s known effect on respiration). We monitored the animal’s core temperature with the device’s bead thermistor probe and, independently, with a RET-4 rectal probe (Physitemp) inserted in parallel with the bead thermistor probe. Further, we kept a record of the surface temperature of the silicone rubber heating pad directly under the anesthetized mouse. The controller was set for 35°. The data indicate that the core temperature of the mouse varied less than 1.3° over the 6 hours, despite the 15° differential between ambient room temperature (20°) versus the set core temperature (35°) (Fig. 6). Importantly, the core temperature of the mouse remained at 35±0.6° without any supplemental warming device (heat lamp, blanket, etc.). The LiPo battery was changed once over the 6-hour period.

**Figure 6.**
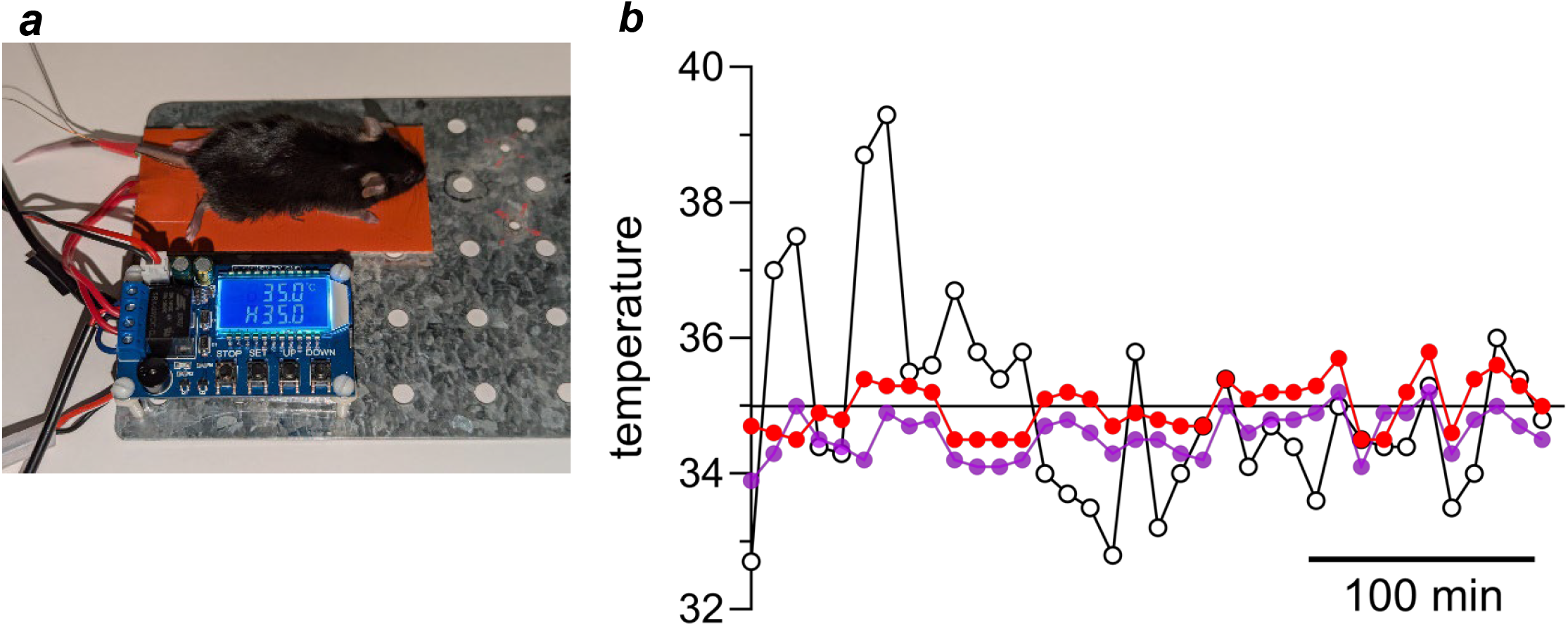
Validation of homeothermic heating pad. An anesthetized mouse was kept on a platform in an open room (ambient temperature, 20°) without any other source of heat for 6 hours. The device was set to maintain the core temperature of the mouse at 35°. ***a***, the experimental set up. ***b***, temperatures recorded by device’s rectal bead thermistor probe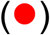; by an independent RET-4 (Physitemp) rectal probe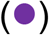; and by a separate temperature probe placed on the silicone heating pad under the mouse (∘).

Second, we have exclusively used the battery-powered warming pad for *in vivo* confocal calcium imaging of sensory ganglion activity [38, 39]. The experiments typically last 6 hours or more with the mouse remaining in a surgical plane of anesthesia for the duration. We have found it important to keep the core temperature of the mouse at 31.5°. Surgical preparations are carried out in a separate room from the imaging rig, requiring transport of the anesthetized mouse fixed on a custom-built stereotaxic device with the warming pad attached for a distance of about 40 m. Ambient room temperatures vary from 20° to 24°. The core temperature of the mouse as measured by the device’s rectal temperature probe (bead thermistor) has consistently remained constant and the animal kept in stable physiological condition throughout the entire 6+ hours without any auxiliary heating. Neural activity has remained robust (Fig. 7), validating the reliability of the battery-powered warming pad to keep the animal normothermic.

**Figure 7.**
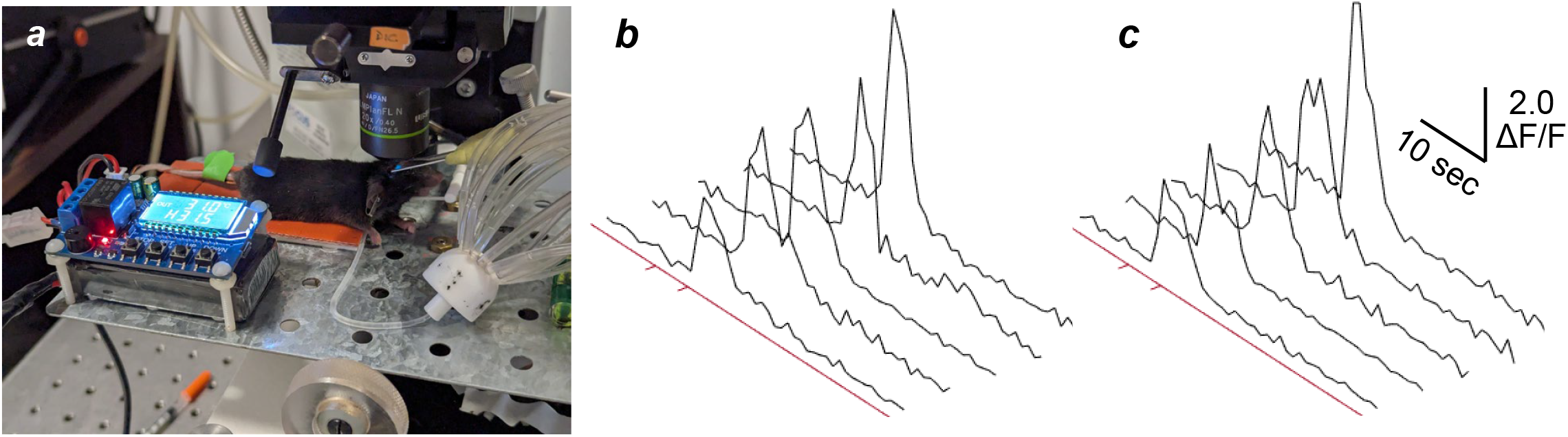
Warming pad during *in vivo* imaging of neuronal activity keeps an anesthetized mouse at 31.5° and allows robust, stable recording for several hours. ***a***, view showing mouse placed on warming pad, its head stabilized with custom-made holder under 20X objective of a confocal microscope. A scanning laser (488 nm) illuminates the geniculate ganglion and images taste-evoked activity in gustatory sensory neurons that express the calcium-sensitive reporter, GCaMP6s. Large white manifold (right) guides taste stimuli into the oral cavity. See [38, 39] for details. ***b***, examples of geniculate ganglion neuron responses (change in GCaMP6s fluorescence, ΔF/F) to a 5 sec presentation of 300 mM sucrose solution into the oral cavity, imaged at beginning of experiment. Ticks on red line indicate time of stimulus delivery. ***c***, examples of responses from 5 different ganglion neurons, recorded after the mouse had been anesthetized and maintained on the warming pad in a surgical plane of anesthesia for 6 hours. The amplitudes, durations, and latencies of the taste-evoked responses are remarkably similar to those recorded several hours earlier.

These tests validate the homeothermic controller and firmly establish the ability of the heating pad device to maintain an anesthetized rodent at a constant core temperature for long durations.

## 4. Discussion

This report describes a small, portable, battery-powered homeothermic heating pad for maintaining normothermia in anesthetized mice and rats. This device was shown to keep the core body temperature of a deeply anesthetized mouse stable over several hours without any supplemental heating source, despite a large differential between room temperature and core body temperature. We illustrated the utility of this heating pad for confocal *in vivo* imaging of cranial ganglion activity in mice.

All the components for the device are available online, for example from *Amazon*. The total cost for all components is less than $100. Commonly used tools are used to construct the unit. No specialized skills, apart from rudimentary soldering, are required to build the device. The warming pad is powered by a high-capacity LiPo battery, commonly used for drones. Finally, the entire device can be assembled in less than 30 minutes when the parts are laid out as described.

This small battery-powered homeothermic heating pad is especially useful for experimental situations where anesthetized animals are prepared at one site (such as a surgical suite) and transported to another (such as an imaging rig) for experimental procedures. Because the device is battery-powered, there are no power cables to disconnect and no separate thermocontroller unit to move. Moreover, being battery-powered, it does not generate electrical noise for electrophysiological recordings, a nagging problem with commercial devices powered by 120/240V. Finally, the heating pad can be readily adapted to custom-built stereotaxic frames, holding boxes, animal cages, or other situations.

In short, the homeothermic battery-powered heating pad described here will be of great utility for studies that use anesthetized mice or rats.

## CRediT authorship contribution statement

**Isabella Fleites**: Validation, Formal analysis, Investigation, Data Curation, Writing - Review & Editing; **Kevin Morales**: Software, Formal analysis, Writing - Review & Editing; **Stephen Roper**: Conceptualization, Methodology, Software, Validation, Investigation, Resources, Data Curation, Writing - Original Draft, Writing - Review & Editing, Visualization, Supervision, Project administration, Funding acquisition

